# Geometric and Topological Approaches to Shape Variation in *Ginkgo* Leaves

**DOI:** 10.1101/2020.10.23.352476

**Authors:** Haibin Hang, Martin Bauer, Washington Mio, Luke Mander

## Abstract

Leaf shape is a key plant trait that varies enormously. The diversity of leaf shape, and the range of applications for data on this trait, requires frequent methodological developments so that researchers have an up-to-date toolkit with which to quantify leaf shape. We generated a dataset of 468 leaves produced by *Ginkgo biloba*, and 24 fossil leaves produced by evolutionary relatives of extant *Ginkgo*. We quantified the shape of each leaf by developing a geometric method based on elastic curves and a topological method based on persistent homology. Our geometric method indicates that shape variation in our modern sample is dominated by leaf size, furrow depth, and the angle of the two lobes at the base of the leaf that is also related to leaf width. Our topological method indicates that shape variation in our modern sample is dominated by leaf size and furrow depth. We have applied both methods to modern and fossil material: the methods are complementary, identifying similar primary patterns of variation, but also revealing some different aspects of morphological variation. Our topological approach distinguishes long-shoot leaves from short-shoot leaves and both methods indicate that leaf shape influences or is at least related to leaf area.

## Introduction

Leaf shape is a fascinatingly diverse plant trait. It can vary between taxa, between individuals in different populations of the same species, and for some species there are striking variations in leaf shape within a single plant, a phenomenon known as heterophylly. Additionally, different regions of a leaf expand at different rates during development, and this leads to allometric changes in shape as a leaf grows. Leaves are primary sites of photosynthesis and play a central role in the growth and survival of a plant, and work has shown that variation in leaf shape may be related to thermoregulation, the constraints of hydraulics and mechanics, patterns of leaf expansion, as well as the avoidance of herbivory and the optimal interception of light (Nicotra *et al*., 2011). Leaf shape is therefore a trait for which there are many functional trade-offs, and from an ecological perspective may be viewed “not as a single major axis, but rather as an option that fine tunes the leaf to its conditions over both short and evolutionary time spans” (Nicotra *et al*., 2011, p. 547).

The taxonomic and ecological significance of leaf shape has led to the development of numerous methods to characterize this trait. Certain methods rely on largely qualitative observation. For example, aspects of leaf shape can be described using specialist terminology (Leaf Architecture Working Group 1999), which allows leaves to be placed into categories based on their gross morphology, and this approach has proved useful in studies of plant architecture (e.g. Leigh, 1999; Barthelemy & Caraglio, 2007) and studies of fossil leaves that may not be preserved in their entirety (e.g. Johnson, 1992). Other methods for characterising leaf shape are based on morphometric measurements of certain features on a leaf, which can either be made manually by human researchers or computationally using image analysis software. For example, Leigh *et al*. (2011) described leaf shape using measurements of leaf area and leaf dissection (leaf perimeter/area) in the context of plant hydraulics, and Royer *et al*. (2005) used the same measure of leaf dissection to investigate the relationship between mean annual temperature and leaf shape. Measurements of such morphological features are often used to generate indices of leaf shape, such as compactness (perimeter^2^/area) and shape factor (4π x leaf area/perimeter^2^), which are used to summarize aspects of leaf shape and show how it relates to the environment or has changed through time (Royer *et al*., 2008, 2009; Peppe *et al*., 2011; Bacon *et al*., 2013). Morphometric techniques that use landmarks (a constellation of discrete anatomical loci, each described by 2- or 3-dimensional Cartesian coordinates to quantify morphology (Thompson 1942; Bookstein 1996; Webster & Sheets, 2010)) have been employed to capture variation in leaf shape (Weight *et al*., 2008) and have highlighted differing developmental and evolutionary contributions to leaf shape (Chitwood *et al*., 2016), while elliptic Fourier analysis has been used to quantify leaf outlines (McLellan 1993; Chitwood & Otoni 2017). Persistent homology—a topological data analysis method—has also been applied to the problem of quantifying leaf shape (Li *et al*., 2018a,b), and represents a morphometric framework to measure plant form that allows comparison of the morphology of different plant organs such as leaves, roots and stems (Bucksch *et al*., 2017; Li *et al*., 2017).

Owing to the diversity of leaf form—and the range of applications for data on leaf morphology—regular methodological experimentation is required so that researchers have an up-to-date toolkit with which to quantify this plant trait. In this paper, we provide such experimentation through a quantitative study of leaf shape in *Ginkgo biloba* L., an extant gymnosperm. We have selected *Ginkgo* as a study system primarily because of the diversity of leaf shapes that are produced by individual specimens (e.g. Leigh *et al*., 2011) and because of the palaeobotanical importance of fossil *Ginkgo* and its extinct evolutionary relatives. In particular, *Ginkgo* and its relatives were important elements of Earth’s vegetation during the Mesozoic Era (∼250–65 million years ago) and fossil leaves of plants that are evolutionary ancestors of living *Ginkgo* are commonly found in sedimentary rocks. These leaves have been widely used to investigate Earth’s ancient atmospheres and environments using their stomatal indices, carbon isotopic composition and physiognomy (see Sun *et al*., 2003; McElwain & Steinthorsdottir 2017; Bacon *et al*., 2013). Consequently, with a view to demonstrating the applicability of our methods to fossil material our study includes a small number (24) of fossil *Gingko* leaves.

Previous work on *Ginkgo* leaf morphology has shown that this plant is characterized by pronounced heterophylly with different leaf forms borne on long shoots versus short shoots (Critchfield 1970; Dorken 2013). The leaves of long shoots are typically smaller and can have a deep wide furrow and a dissected margin, while the leaves of short shoots are typically larger and can have a less pronounced furrow (Leigh *et al*., 2011). The variability of *Ginkgo* leaf morphology is emphasized by measures of specific leaf area (the ratio of leaf lamina area to leaf lamina dry mass), which indicate that the form of *Ginkgo* leaves varies not only between the long and short shoots of the plant, but also between the trees of different genders (micro-versus megasporangiate), as well as between juvenile and mature portions of a megasporangiate canopy, and also for short shoots bearing seed and adjacent short shoots without seed (Christianson and Niklas 2011, see also Niklas and Christianson 2011). The hydraulic architecture of *Ginkgo* leaves has been quantified (Carvalho *et al*., 2017) and it has been shown that long-shoot and short-shoot *Ginkgo* leaves have different structural and hydraulic properties, probably related to greater hydraulic limitation of long-shoot leaves during leaf expansion (Leigh *et al*., 2011). Aspects of *Ginkgo* leaf shape are demonstrably sensitive to atmospheric composition, and when quantified by shape factor, extant *Ginkgo* leaves that have been subject to elevated atmospheric SO2 levels in controlled environment chambers are significantly rounder than control leaves (Bacon *et al*., 2013). Additionally, *Ginkgo* leaf shape may also be sensitive to elevation, although the positive relationship between the length:width ratio of leaves and elevation is weak (Xie *et al*., 2013).

Our study builds on this body of previous work by taking an exploratory approach to the morphology of *Ginkgo* leaves. We do not initially focus on any specific morphological features such as leaf length or the nature of the leaf margin, but instead use geometric and topological methods to reveal the features that explain the observed variation in leaf shape. Our overall goal is to provide an illustration of how these methods can be applied to the problem of quantifying leaf shape, and our specific aims are as follows: (1) to develop a geometric method and a topological method for quantifying leaf shape; (2) to apply these methods to the leaves of living *Ginkgo* in order to reveal which features explain the observed variation in the shape of sampled leaves; (3) to compare the results produced by the two methods in order to explore the degree to which they reveal different aspects of morphological variation; and (4) to apply our methods to fossil leaves of ancient evolutionary relatives of living *Ginkgo* in order to confront a degree of morphological variation not present in our sample of living *Ginkgo*, and to demonstrate how they could be used to study the evolution of leaf shape through geological time.

### A Dataset of Modern and Fossil Leaves

Mature and fully expanded leaves were harvested from a reproductively immature *Ginkgo biloba* tree growing in partial shade as a specimen on the campus of The Open University, UK. The specimen was ascended using a ladder and seven branches growing towards the West at approximately halfway up the specimen were removed from the trunk using a saw. Every leaf growing on each branch was plucked from the base of the petiole and dried in a plant press. A total of 468 leaves from a mixture of short-shoots and long-shoots were collected from the specimen. Each of these leaves was photographed next to a scale bar using a digital camera positioned 20cm above a light box. Twenty-two fossil leaves produced by evolutionary relatives of living *Ginkgo biloba* were extracted from the collections of the Natural History Museum in London, and two fossil leaves were extracted from the geology collections of the School of Environment, Earth and Ecosystem Sciences, The Open University (Table 1). Each fossil leaf was photographed next to a scale bar using a digital camera and the outline of each fossil was traced using Adobe Illustrator to create a digital outline of each leaf. The petioles of fossil leaves are frequently broken, distorted or completely absent as a result of the fossilization process. A central goal of our manuscript is to compare living and fossil *Ginkgo* leaves and in order to facilitate this, we have excluded the petiole from our analyses. Our analyses are therefore focussed on the shape of *Ginkgo* leaf blades. Our dataset of modern and fossil *Ginkgo* leaf images is available in the Supplementary Information.

**Table 1.** Fossil *Ginkgo* leaves housed in the collections of the Natural History Museum, London, and The Open University that we have investigated in this paper.

| Specimen Name | Accession Number | Age | Country | Location | Specimen Number (This Study) |
| --- | --- | --- | --- | --- | --- |
| *Ginkgo cranei* | NHM: V.68763 | Paleocene | United States | North Dakota | fossil_1 |
| *Ginkgo cranei* | NHM: V.68764 | Paleocene | United States | North Dakota | fossil_2 |
| *Ginkgo gardneri* | NHM: V.14834 | Eocene | Scotland | Isle of Mull | fossil_3 |
| *Ginkgo gardneri* | NHM: V.14838 | Paleocene/Eocene | Scotland | Isle of Mull | fossil_4 |
| *Ginkgo gardneri* | NHM: V.18436 | Eocene | Scotland | Isle of Mull | fossil_5 |
| *Ginkgo gardneri* | NHM: V.24999 | Eocene | Scotland | Isle of Mull | fossil_6 |
| *Ginkgo sp.* | NHM: V.2477 | Eocene | Scotland | Isle of Mull | fossil_7 |
| *Ginkgo digitata* | NHM: V.24587 | Cretaceous | Australia | Queensland | fossil_8 |
| *Ginkgo digitata* | NHM: V.39211 | Jurassic | England | Yorkshire | fossil_9 |
| *Ginkgo digitata* | NHM: V.13503 | Jurassic | England | Yorkshire | fossil_10 |
| *Ginkgo digitata* | NHM: V.10316 | Jurassic | England | Yorkshire | fossil_11 |
| *Ginkgo huttonii* | NHM: V.60195 | Jurassic | England | Yorkshire | fossil_12 |
| *Ginkgo huttonii* | NHM: V.3580 | Jurassic | England | Yorkshire | fossil_13 |
| *Ginkgo huttonii* | NHM: V.40511 | Jurassic | England | Yorkshire | fossil_14 |
| *Ginkgo huttonii* | NHM: V.39210 | Jurassic | England | Yorkshire | fossil_15 |
| *Ginkgo huttonii* | NHM: V.978 | Jurassic | England | Yorkshire | fossil_16 |
| *Ginkgo huttonii* | NHM: V.979 | Jurassic | England | Yorkshire | fossil_17 |
| *Ginkgo longifolius* | NHM: V.39209 | Jurassic | England | Yorkshire | fossil_18 |
| *Ginkgo siberica* | NHM: V.58618 | Jurassic | England | Yorkshire | fossil_19 |
| *Ginkgo digitata* | NHM: V.3423 | Jurassic | England | Gloucestershire | fossil_20 |
| *Ginkgo digitata* | NHM: V.3429 | Jurassic | England | Gloucestershire | fossil_21 |
| *Ginkgo siberica* | NHM: V.19238 | Jurassic | Russia | Irkutsk | fossil_22 |
| *Ginkgo huttonii* | Open University geology collection | Jurassic | England | Yorkshire | fossil_23 |
| *Ginkgo huttonii* | Open University geology collection | Jurassic | England | Yorkshire | fossil_24 |

### A Geometric Approach to Quantifying the Shape of Leaves

#### Methods

Building on previous work using elastic curves to quantify leaf shape (Laga *et el*. 2012, 2014), we represented each *Ginkgo* leaf blade by its boundary curve, with values mapped in the plane (two dimensional Euclidean space) (Fig. 1). When considering these representations of *Ginkgo* leaves we factored out the actions of rotation and translation and reparameterization. For example, two identical leaves could each be represented by their boundary curves, but each curve could be considered distinct from one another if they differed only by rotation (a curve could be presented at 90 degrees on top of the other for instance), but our analysis factors out such actions. It is possible to also factor out the action of scaling and we do this in an analysis of leaf shape versus leaf area.

**Fig. 1.**
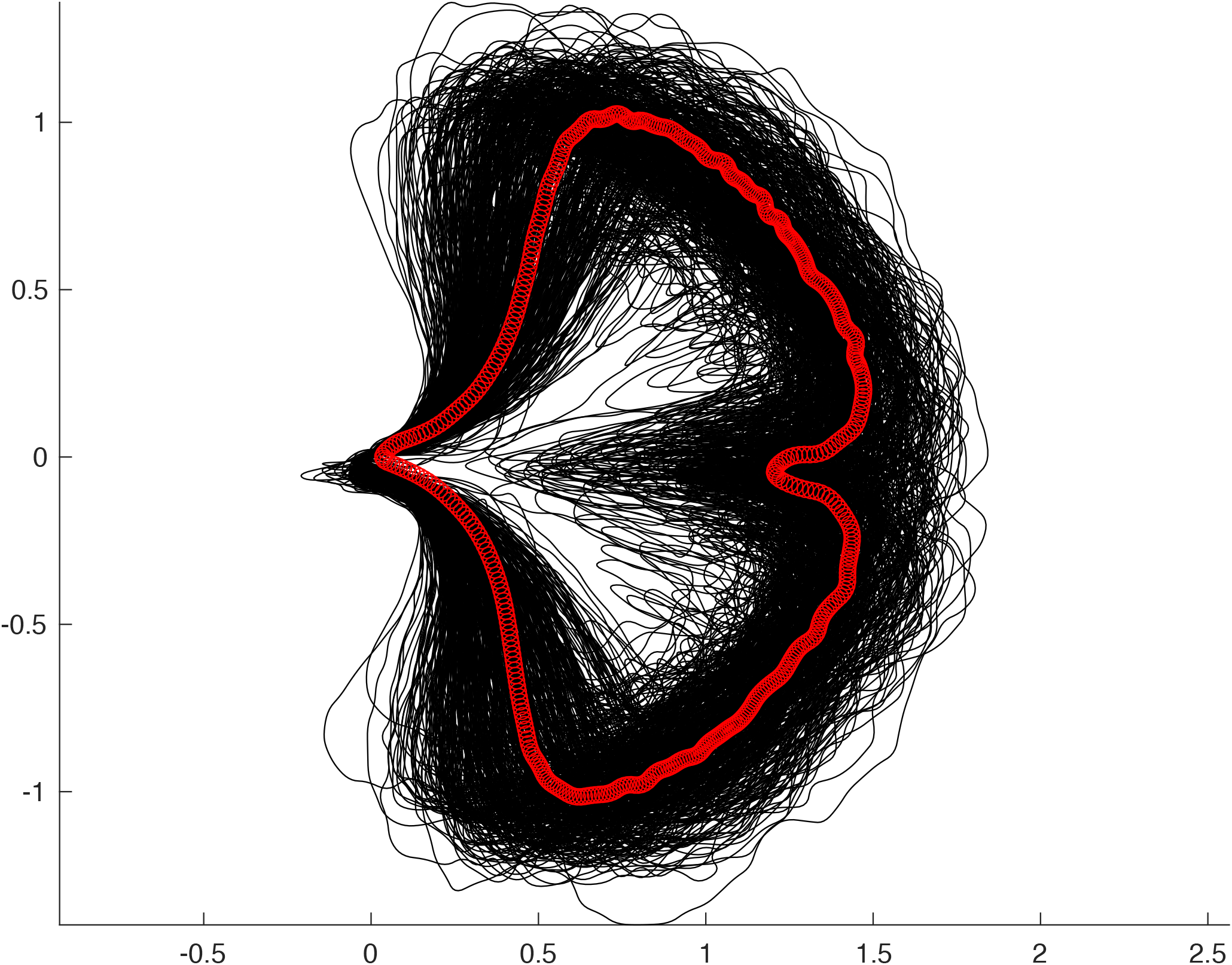
Collection of all 468 *Ginkgo biloba* leaves in our dataset represented by their boundary curves (black lines) with the Karcher mean leaf shape superimposed (red line).

To quantitatively model morphological variation in our sample of *Ginkgo* leaves, we introduce a similarity measure for shapes that serves as the basis of statistical analysis. This is an intricate process for two main reasons: (1) the infinite dimensionality of the ensemble of all shapes; and (2) the non-linearity of shape space. To overcome this difficulty, we appeal to the concepts of Riemannian geometry, and use a Riemannian metric that quantifies the difficulty of morphing one boundary curve onto another by measuring the geodesic distance between the curves, accounting for rotations, translations and reparameterizations. This enables us to quantify shape similarity as the minimal deformation cost to reshape a curve, in this case a *Ginkgo* leaf contour. Despite the nonlinear nature of shape space, this framework allows us to calculate mean shapes and locally linearize shape data about the mean, which, in turn, lets us employ standard statistical methods on linearized data to analyse the shape variation present in our sample of *Ginkgo* leaves.

The Riemannian metric we employ is grounded on principles of linear elasticity and is formally defined on the ensemble of parametric curves, but its invariance properties ensure that it descends to a shape metric. A precise definition of the metric and a discussion of its main properties may be found in Bauer *et al*. (2017, 2019) (see also Klassen *et al*. (2004) for related shape metrics). In practice, the comparison of *Ginkgo* leaf boundary curves is a shape-matching problem, and to solve we discretized the boundary curve of each leaf using a finite dimensional representation. This reduces the problem of comparing leaf boundary curves to a finite-dimensional optimization problem that can be solved with standard methods of numerical optimization. We use principal component analysis (PCA) to uncover the principal modes of shape variation in *Ginkgo* leaves.

The essential steps in this approach are: (1) image processing to isolate each leaf from the image background and remove the petiole; (2) find the boundary curve of each leaf blade; (3) discretize boundary curve of each leaf blade using a finite dimensional representation; (4) for each leaf blade, calculate an elastic metric that quantifies the difficulty of morphing one leaf boundary curve onto another; (5) compare leaves and visualize the dominant modes of shape variation among leaves using PCA. The code underlying our geometric approach is fully open source available at https://github.com/h2metrics/h2metrics. The provided package contains detailed documentation and user-friendly working examples.

#### Results

We calculated the Karcher mean of our sample of modern *Ginkgo* leaves (Fig. 1) and then locally linearized the data about the mean in order to uncover the principal modes of leaf shape variation. This was accomplished by solving a shape-matching problem between the mean and each leaf in the dataset.

Principal component analysis on the linearized data indicated that approximately 30 components are needed to explain 80% of the shape variation in our sample of *Ginkgo* leaves (Fig. 2a), and we graphically display the principal modes of leaf shape variation using geodesic PCA plots (Fig. 2b–d). The first mode is predominantly leaf size (first principal component, Fig. 2b), the second mode relates to the nature of the leaf margin (second principal component, Fig. 2c), and the third and fourth modes are the depth of the furrow that separates the two lobes of the typical *Ginkgo* leaf, together with the angle of the two lobes at the base of the leaf that is also related to leaf width (third principal component, Fig. 2d). Some leaves, for example, have a very deep furrow whereas others have no furrow at all. Similarly, some leaves have lobes that are quite pointed and curve backwards towards the leaf base, whereas others have lobes that do not curve backwards.

**Fig. 2.**
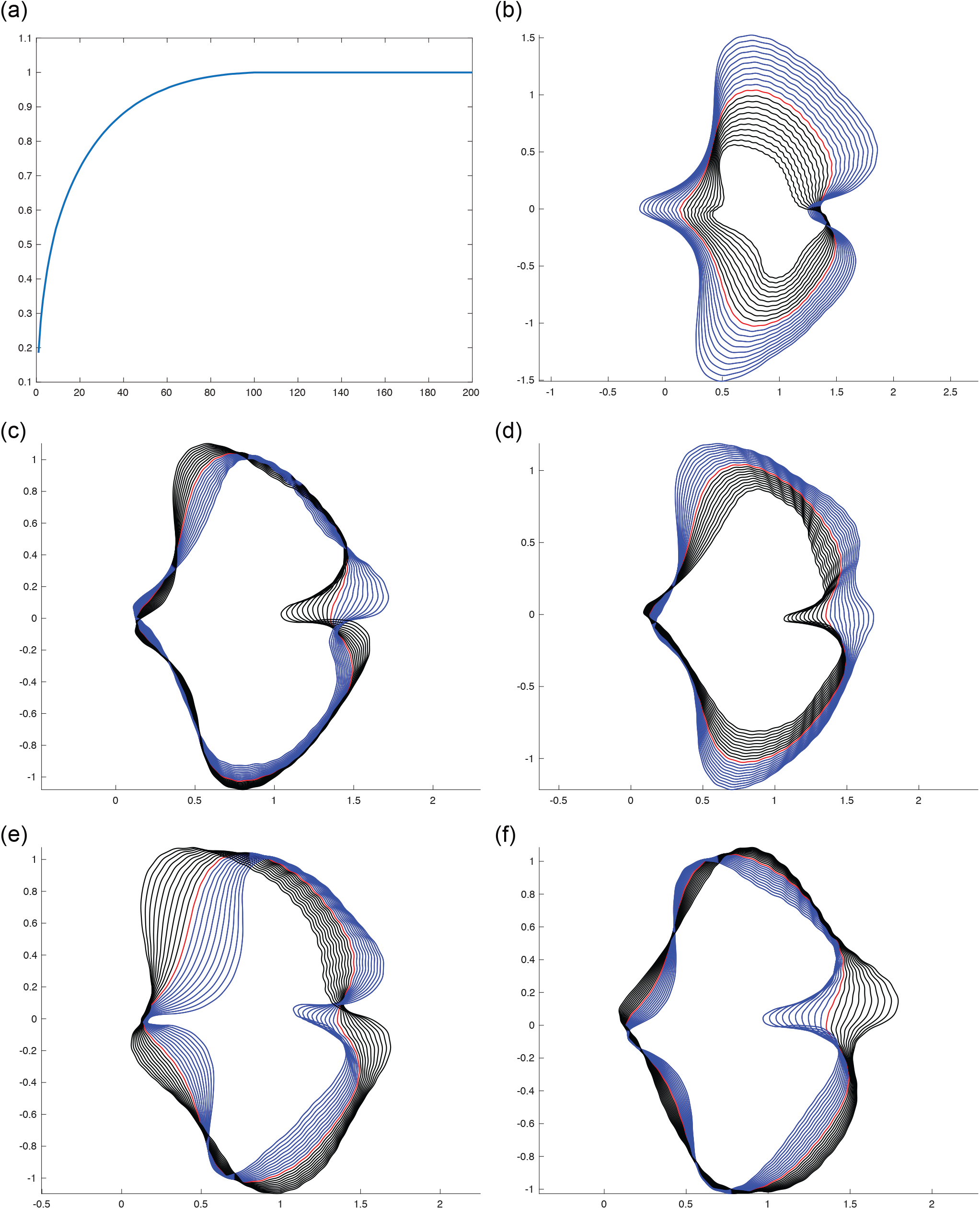
Geodesic PCA plots of *Ginkgo* leaves represented in the tangent space of the mean. Variance explained by all components (a), the first principle component (b), second principle component (c) and third principle component (d). Analysis with scaling factored out, first principle component (e) and second principle component (f).

Examples of variability in terms of the morphological features identified by our geodesic plots (Fig. 2b–d) can be seen in a PCA ordination of our dataset of *Ginkgo* leaves (Fig. 3a). Leaves towards the left are relatively small and leaves towards the right are relatively large (Fig. 3a). Leaves to the bottom are typically more dissected and have a relatively deep furrow, whereas leaves to the top are typically less dissected and have a relatively shallow furrow (Fig. 3a). This plot also highlights that the morphological space occupied by our sample of *Ginkgo* leaves, as delineated by our geometric approach, is organized as a continuous distribution of datapoints without separate clusters. Most data points are concentrated towards the center of the ordination, and the distribution of data points becomes sparser with increasing distance from the center (Fig. 3a).

**Fig. 3.**
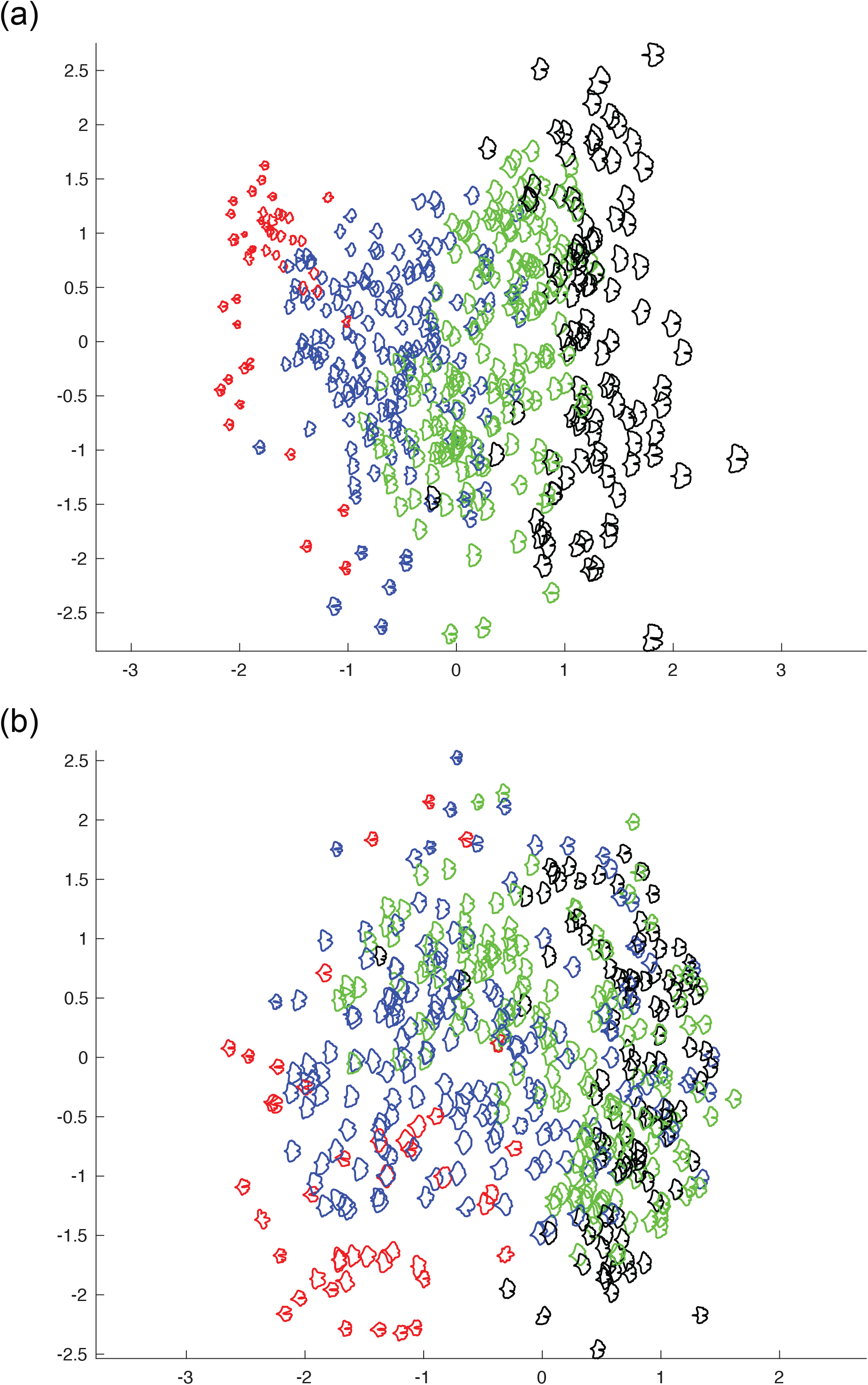
PCA ordination scatterplot (PC1 on horizontal axis, PC2 on vertical axis) showing the morphological variation among 468 modern *Ginkgo* leaves that is revealed by our geometric approach to leaf shape incorporating scale (a), and with scale factored out (b). Leaf area groups based on their areas: area ≤ 8 (red); 8 < area ≤ 16 (blue); 16 < area ≤ 16 (green), area ≥ 24 (black); area in cm^2^.

### A Topological Approach to Quantifying the Shape of Leaves

#### Methods

We employed the topological data analysis technique persistent homology (PH) (Edelsbrunner & Harer 2010; Otter *et al*. 2017; Li *et al*. 2018a,b) and represented each *Ginkgo* leaf in our dataset with a persistence barcode. To construct this barcode, for each point on the contour of a leaf (its boundary curve), we calculated the distance to the point *P* where the leaf blade meets the petiole (Fig. 4a). Distance was measured in pixels and in our source images 152 pixels = 1 cm. All images were downscaled by 1/8 and so 19 pixels = 1 cm in our analyses. For each *r* > 0, we counted the number of connected components formed by the points on the contour whose distance to *P* is greater or equal to *r* and recorded this count as a barcode. For example, for *r* = 8.6, there are 4 connected components (these are the uninterrupted segments of the leaf blade contour, Fig. 4a), so there are *b* = 4 bars over that value of *r* (Fig. 4b). Similarly, for *r* = 7.0, 5.4, 3.8, (Fig. 4a) the corresponding number of bars is b = 3, 2, 1 (Fig. 4b). The barcode summarizes the count as we gradually lower the threshold r, with bars disappearing as connected components coalesce and bars appearing as new components emerge. The coalescence of two connected components follows the elder rule: the first-born bar survives while the younger bar dies. Through this construct, we mapped the dataset of leaves to a dataset of barcodes, with each leaf described by a barcode. In order to facilitate statistical analysis, we vectorized each barcode by listing the length of the bars in decreasing order. Since different leaves may produce barcodes with different number of bars, we padded the tails of the vectors with zeros to make all vectors the same length. In our analysis of modern leaves, statistical analyses were performed on these padded vectors. In our analysis of modern and fossil *Ginkgo* leaves combined, statistical analyses were performed on vectors that were normalized by the length of the first bar (the first component of each normalized vector was therefore 1 and discarded). The essential steps in this approach are: (1) image processing to isolate each leaf from the image background and remove the petiole; (2) find the contour curve of each leaf blade; (3) reparameterize the contour curve of each leaf blade; (4) calculate the persistent homology of each leaf blade; (5) construct the persistence barcode of each leaf blade; (6) compare leaves and visualize the dominant modes of shape variation among leaves using multi-dimensional scaling. An example of the code underlying our topological approach is available at https://github.com/Haibin9632/TDA-of-ginkgo-leaves. The provided code contains documentation and user-friendly working examples.

**Fig. 4.**
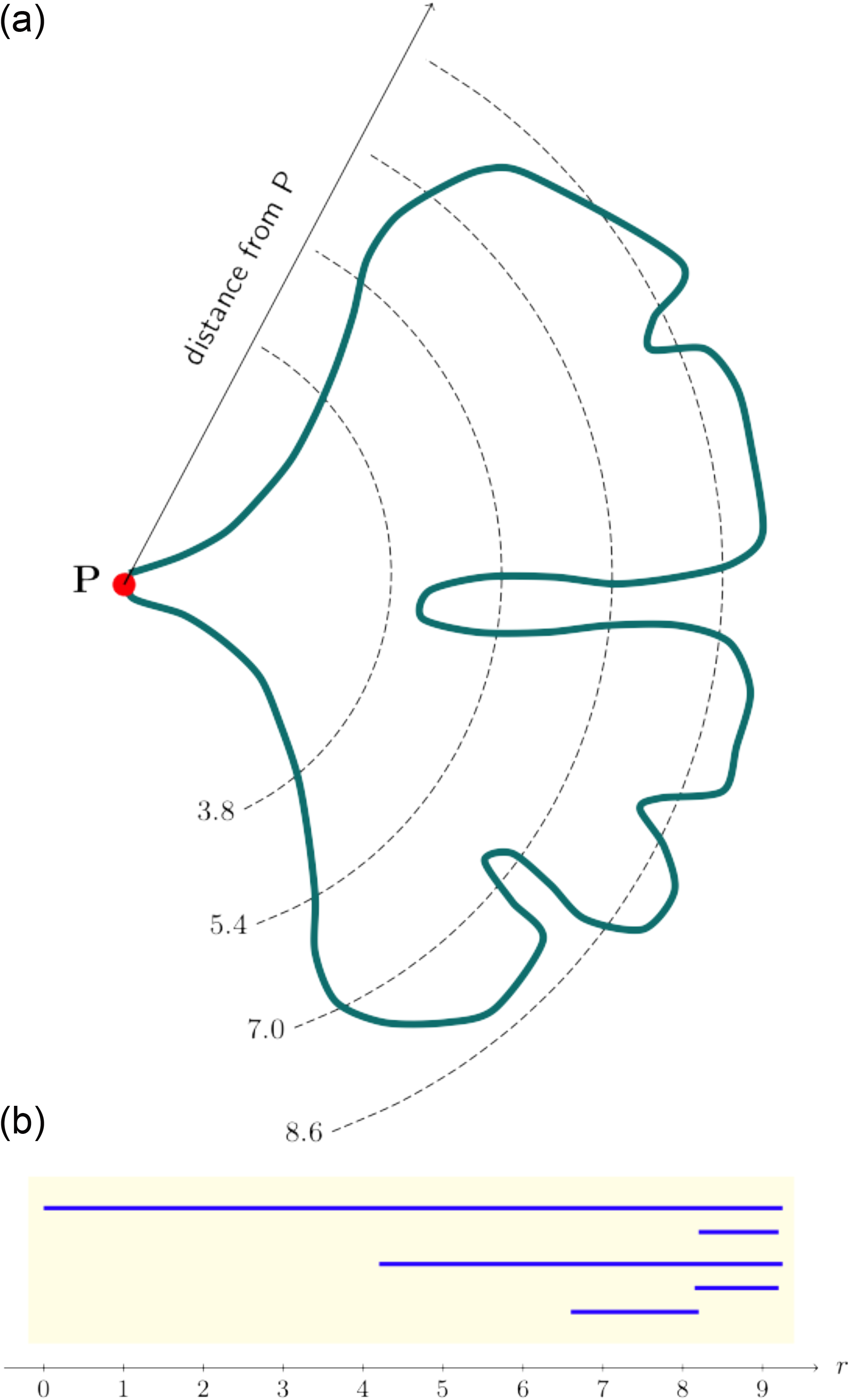
Schematic example showing the construction of a persistence barcode that describes the shape of a *Ginkgo* leaf. Four distances from the point *P* where the leaf blade meets the petiole are shown: *r* = 8.6, 7.0, 5.4, 3.8 (a). At the distance *r =* 8.6, there are four connected components outside the dashed line (a). At the distance *r* = 7.0, there are three connected components, at *r* = 5.4 there are two (the two lobes of the typical *Ginkgo* leaf), while at *r* = 3.8 there is one uninterrupted segment of the leaf blade contour outside the dashed line (a). To construct a barcode that represents a leaf, we do not count the number of connected components at widely spaced intervals as shown in (a). Instead, we perform a count for each *r* > 0, and record the number of connected components as *r* is gradually lowered in a barcode (b).

#### Results

Figure 5 shows the results of PCA applied to the vectorized barcode data. The first PC explains approximately 75% of the total variance and inspection of the PC loadings indicates that it is dominated by leaf length, followed by furrow depth. The second PC explains about 22% of the total variance mainly as variation in the depth of the furrow, followed by (negative) variation in leaf length. This ordination indicates that the morphological space occupied by our sample of *Ginkgo* leaves, as delineated by our topological approach, is organized as a continuous distribution of datapoints without separate clusters, although the majority of leaves lie in the quadrant of PC1 scores –20 to 20 and PC2 scores 0 to –20, and the leaves with PC1 scores < 0 and PC2 scores > 15 are perhaps separated from the other leaves in our sample (Fig. 5). To facilitate visualization of shape variation among our sample of *Ginkgo* leaves, the original leaf images corresponding to two discrete paths, nearly parallel to the first two principal PC axes, are highlighted in Fig. 5. These two paths show contrasting behaviour: PC1 captures a pattern in which larger leaves have a deeper furrow, whereas PC2 captures a pattern in which smaller leaves have a deeper furrow.

**Fig. 5.**
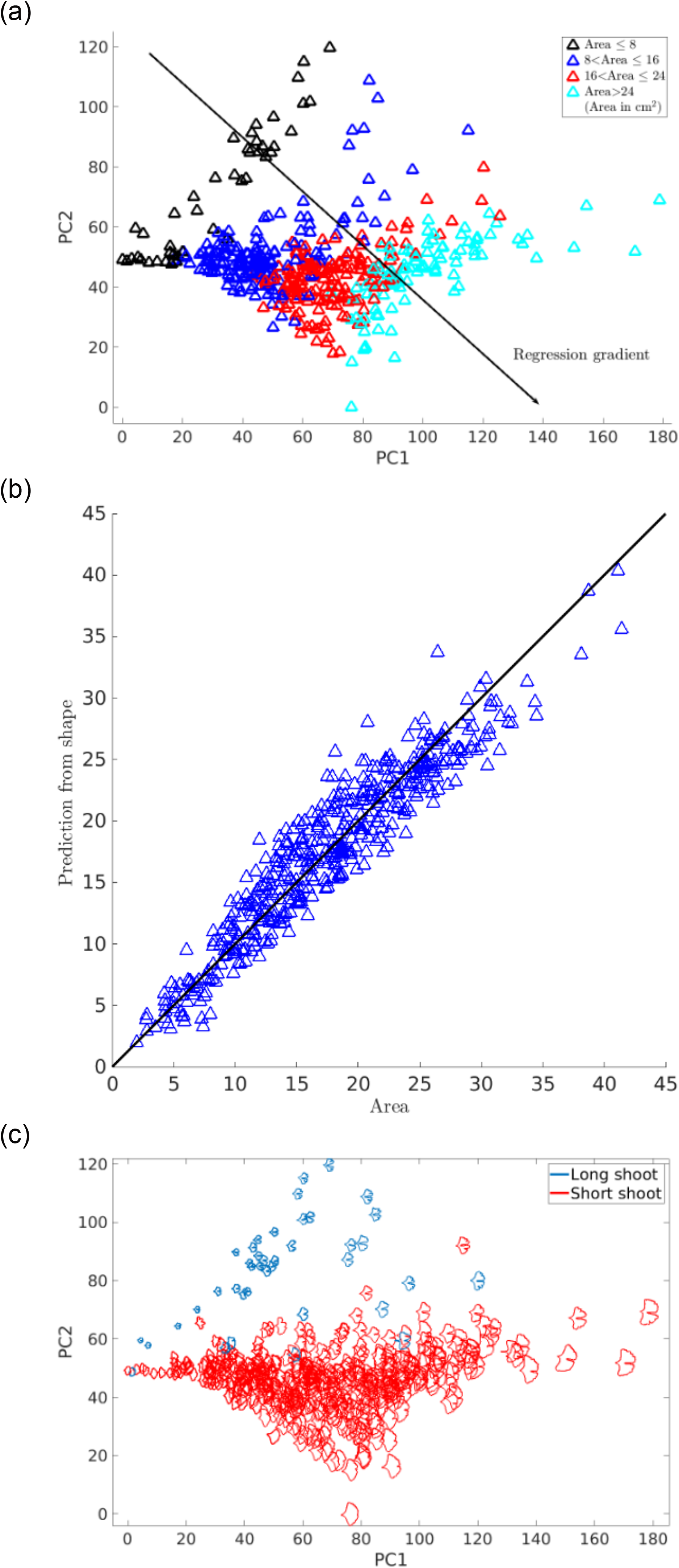
PCA ordination scatterplot showing the morphological variation among 468 modern *Ginkgo* leaves that is revealed by our topological (PH) approach to leaf shape, with datapoints coloured according to four leaf area groups (a). Comparison of leaf area predicted from the first two principal components and measured leaf area; the vertical offset between datapoints and the solid black diagonal 1:1 line of equality indicates the discrepancy between the predicted and measured areas (b). PCA ordination scatterplot with long-shoot and short-shoot *Ginkgo* leaves highlighted (c).

### Application to Fossil *Ginkgo* Leaves

Visual inspection of fossil leaf boundary curves highlights that the diversity of leaf shapes in our collection of *Ginkgo* fossils is greater than that found in our sample of modern *Ginkgo* leaves (compare Fig. 1 and Fig. 6a, see also the Supplementary Information). In particular, several fossil leaves are characterised by multiple deep furrows so that leaf blades consist of multiple lobes rather than just two as in the typical *Ginkgo biloba* leaf, while other fossils have highly dissected leaf margins. This greater diversity in fossil leaf shapes is picked up by both the geometric and the topological approaches we have described, and both indicate that there are fossil leaves situated outside the total range of morphological space occupied by modern *Ginkgo* leaves (Fig. 6b,c). Both approaches also highlight that there are some fossils leaves that are very similar to modern *Ginkgo* leaves, and there are some fossil and modern leaves that overlap in morphological space (Fig. 6b,c).

**Fig. 6.**
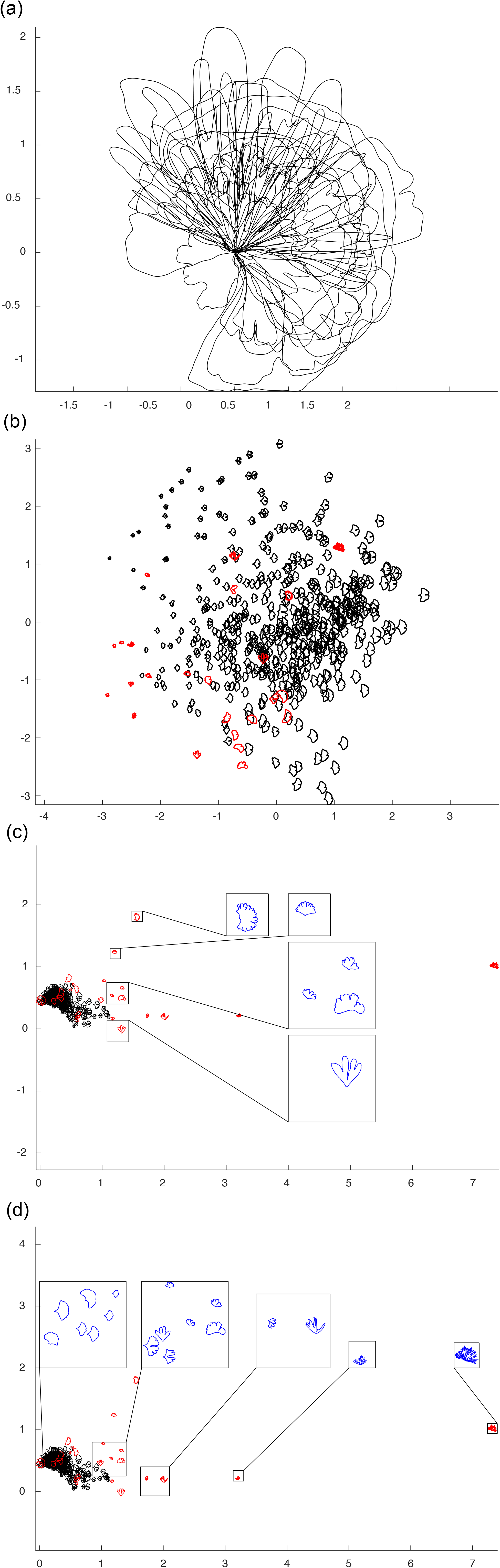
Collection of 24 fossil *Ginkgo* leaves, each represented by their boundary curves (a). PCA ordination scatterplot (PC1 on horizontal axis, PC2 on vertical axis) showing morphological variation of modern *Ginkgo* leaves (black datapoints) together with fossil *Ginkgo* leaves (red datapoints) based on our geometric approach, the PCs together explain 64% of the variation (b). MDS ordination showing morphological variation of modern *Ginkgo* leaves together with fossil *Ginkgo* leaves based on our topological approach, with a vertical transect of enlarged leaves highlighted in blue (c) and a horizontal transect of enlarged leaves highlighted in blue (d). Modern leaves displayed with black datapoints and fossil leaves displayed with red datapoints (b–d).

However, there are differences in the degree to which modern and fossil leaves are separated in morphological space using our two approaches. Using our geometric approach, relatively small leaves with shapes characterised by multiple lobes lie outside the morphological space occupied by modern *Ginkgo* leaves, while relatively large leaves with highly dissected margins plot within the space occupied by modern leaves (Fig. 6b). In contrast, using our topological approach, both of these types of fossil leaves plot outside the morphological space occupied by modern *Ginkgo* leaves (Fig. 6c). Our topological approach very clearly captures similarities and differences between modern and fossil leaves that are expected on the basis of their visual appearance alone (Fig. 6c), whereas using our geometric approach the distinction between modern and fossil leaves is not as clear (Fig. 6b).

## Discussion

### Comparison of Approaches

The two approaches we have described in this paper measure leaf shape in different ways: our geometric approach is based on analysing boundary curves with an elastic metric (Fig. 2), whereas our topological approach is based on measuring the number of connected components as a leaf is partitioned into different segments (Fig. 4). Despite these differences, the two approaches both indicate that leaf size and the nature of the furrow separating the two lobes of a typical *Ginkgo* leaf are primary features that explain the observed variation in leaf shape, and both approaches also distinguish the leaves of *Ginkgo* long shoots from those of short shoots. In the PCA summary of our geometric approach, the long shoot leaves are situated to the top left of the plot with low PC1 scores and high PC2 scores, and form a sparsely occupied region of morphological space (Fig. 3). In the PCA summary of our topological approach, the long shoot leaves are situated in the top left of the plot with low PC1 scores and high PC2 scores, and form a sparsely occupied region of *Ginkgo* leaf morphospace (Fig. 5).

There are also certain differences in the morphological features pinpointed by each approach. For example, our geometric approach suggests that the angle of the two lobes at the base of the leaf (also related to leaf width) is an important mode of morphological variation in the population of leaves we have studied (Fig. 2c), but this aspect of leaf morphology is not clearly picked up by our topological approach (Fig. 5). Additionally, our topological approach is able to quantify the nature of the indentations in the leaf margin more clearly than our geometric approach. This is because our topological features, by design, precisely measure the depth of indentations—from large furrows to minor crenulations—in the leaf margin. The vectors we used in our topological analysis of modern and fossil *Ginkgo* leaves were normalized by the length of the first bar, and each vector therefore encodes the depths of the various indentations in the leaf margin relative to absolute leaf size ordered from deep to shallow. This is highlighted in the horizontal transect in Figure 6d: to the left are modern and fossil *Ginkgo* leaves that lack indentations, whereas to the right are leaves with increasingly complex indentations, but the size of each leaf in each highlighted group varies considerably. In the language of descriptive botany, the MDS axes highlight types of leaf dissection, with axis one representing a gradient from no dissection (low axis one scores) to many relatively deep indentations (high axis one score) (Fig. 6d), and axis two representing a gradient from few relatively deep indentations (low axis two scores) to many relatively shallow indentations (high axis two scores) (Fig. 6c). This morphological feature may only be recorded in the higher orders of variation in our geometric approach (fourth and fifth principal components for our modern *Ginkgo* leaves, see Fig. 2e,f). The two approaches we have described are therefore complementary, identifying similar primary patterns of variation, but also revealing some different aspects of morphological variation.

From the perspective of PH applied to the problem of quantifying leaf shape, previous approaches have been based on measurements of the Euler characteristic curve (Li *et al*., 2018a,b). Our approach is different in that we have constructed a persistence barcode from a count of connected components formed by points on a contour at incremental distances from the base of a leaf blade (Fig. 4), and this demonstrates an alternative means by which PH can quantify leaf shape. Oftentimes, a challenge in the use of PH is the interpretation of a persistence barcode (e.g. Otter *et al*., 2017), but for the barcodes we have generated here, the length of the longest bar represents the largest distance to *P* (Fig. 4) and is therefore a quantifier of leaf size, while the next longest bar relates to the depth of the furrow in a *Ginkgo* leaf that displays this trait, and other smaller bars relate to the depth of smaller indentations in the leaf margin. The statistical interpretation of persistence barcodes is also challenging, and as noted by Otter *et al*. (2017, p. 3) for example, “the space of barcodes lacks geometric properties that would make it easy to define basic concepts such as mean, median, and so on”. In contrast, the framework of our geometric approach allows for the calculation of mean shapes and the linearization of data around the mean, and this highlights the complementary nature of the two approaches to leaf shape we have described in this paper.

### *Leaf Shape Versus Leaf Area: the Physiognomy of* Ginkgo *leaves*

Variation in leaf area is first-order mode of variation in our dataset of *Ginkgo* leaves, and experimental work on extant *Ginkgo* leaves has shown a coordinated area–shape response to elevated atmospheric SO_2_ levels. In particular, leaves grown in a high SO_2_ atmosphere were both rounder and smaller than control leaves (Bacon *et al*., 2013). To investigate the relationship between the area and shape of *Ginkgo* leaves in more detail we measured the area of each leaf blade using line integrals and then partitioned the leaves into four groups based on their areas (area ≤ 8, 8 < area ≤ 16, 16 < area ≤ 16, area ≥ 24; area in cm^2^). We then displayed how leaf area varies across morphological space by colour-coding each of these four groups in our PCA ordinations. The colour coding highlights that for our geometric approach variation in leaf area occurs primarily along the first principal component so that leaves increase in area as their size increases (Fig. 3a). In our topological approach variation in leaf area occurs along both the first and second principal components—the boundaries between leaf area groups are orientated obliquely—indicating that leaves increase in area as their size increases and their furrows get shallower (Fig 5a).

For our geometric approach we investigated the relationship between leaf area and leaf shape further by undertaking an analysis of *Ginkgo* leaf blade shape in which we factored out the effects of scaling in the comparison of leaf boundary curves (two leaves were not considered distinct if they only differed in their area). Geodesic PCA plots show the first two principal components of our scale-invariant analysis, and show that the first principal component relates primarily to the shape of the leaf blade at its base (Fig. 2e) and the second principal component relates to the depth and width of the furrow (Fig. 2f). In a PCA ordination summarizing this analysis, leaves that have essentially the same boundary curve but belong to different area groups plot very close to each other in morphological space, and the gradient between leaf area groups is less clear (Fig. 3b); both results are expected since leaf area was factored out in this analysis. However, although the boundaries between area groups is less clear, there remains a trend from leaves with a small area in the lower left of the ordination to leaves with a large area to the right of the ordination, and there is minimal overlap between leaves with an area <8cm^2^ and leaves with an area >24cm^2^ (Fig 3b).

For our topological approach we investigated the relationship between leaf area and leaf shape further by analysing how accurately leaf area can be inferred from just the two dominant principal components of shape derived from our topological representation of *Ginkgo* leaves. We estimated leaf area by performing a linear regression of leaf area over the first two principal components (the regression gradient points diagonally to the lower right of the PCA ordination (Fig. 5a)), and then compared this estimated area to the measured area of each leaf (Fig. 5b). We then quantified the discrepancy between predicted and measured areas by calculating an R^2^ statistic that shows 90% of the variation in the estimated leaf area is explained by variation in measured leaf area and *Ginkgo* leaf shape, as represented by our topological (PH) approach, is therefore strongly related to leaf area.

Taken together, our analyses of leaf shape versus leaf area suggest three things. Firstly, datapoints from different area groups overlap in PCA summaries of both our geometric and topological representation of *Ginkgo* leaves (Fig. 3a and Fig. 5a) and this highlights that while clearly an important mode of morphological variation, leaf area is not the only trait responsible for organising the distribution of datapoints in *Ginkgo* leaf morphological space. This is emphasized by the offset between measured leaf areas and those predicted using our topological representation (Fig. 5b). Secondly, the weak leaf area trend that is present in our geometric scale-free analysis (Fig. 3b), suggests that leaf area may itself exert an influence on the other morphological traits such as furrow depth and the nature of the leaf margin. These observations may support the idea (based on work with angiosperm leaves rather than gymnosperm leaves) that leaf shape may “a trait for which there are many quite varies functional trade-offs” and that may be an “option that fine tunes the leaf to its conditions” (Nicotra *et al*. 2011; p. 547). Finally, our observations support reports indicating that leaf shape is at least correlated with leaf area in *Ginkgo* (Bacon *et al*. 2013; see also Lin *et al*. 2020 for an area–shape correlation among bamboo leaves).

### Image Segmentation

Image segmentation—the partitioning of a digital image into multiple segments—is a key step in any study involving the computational analysis of digital imagery. In this study, the goal of image segmentation was to represent each leaf by its outline. For our sample of modern *Ginkgo* leaves we were able to achieve segmentation computationally because the leaves themselves were whole, free from damage such as indentations in the leaf margin, and the images were free from major defects such as blurring. However, for the fossil *Ginkgo* leaves we have analysed, segmentation involved tracing the outline of each fossil leaf by hand rather than delineating the leaf margin computationally. In some cases of damage to a specimen, the original undamaged margin of a leaf was extremely faint, sometimes only visible using a microscope, whereas in others the leaf margin was interrupted by a scratch or hidden by a small piece of sediment (see the Supplementary Information). In situations such as these, knowledge of the processes leading to the formation and preservation of fossil leaves was used to calibrate a restoration of the fossil outline to what was judged to be its original state. This process introduces a source of potential error that is not quantified, and future work could explore how to automate elements of this image segmentation step, perhaps using a library of fossil leaf outlines produced by manual tracing to train a classifier, or perhaps repairing defects in the leaf margin computationally using techniques from inpainting (see Bertalmio *et al*., 2000). The latter could be particularly valuable in studies of leaves where damage by insects is high such as in lowland moist tropical rainforests.

Discussion of image segmentation is important because it can be a factor that limits the scope of studies that rely on the computational analysis of biological imagery. In particular, we feel that image segmentation will become a key issue if methods such as those we have described here are to be upscaled and automated to analyse large numbers of fossil leaves palaeoecologically.

### Future Applications

The inclusion of fossil leaves in this exploratory analysis (Fig. 6) indicates that both the PH framework and geometric methods based on elastic curves have potential application to evolutionary and palaeoecological problems that require data on leaf shape in the geological past (e.g. Johnson, 1992; Leaf Architecture Working Group, 1999; Royer *et al*., 2008, 2009; Peppe *et al*. 2011; Bacon *et al*., 2013). Shape data derived from these approaches could also be used as classifiers in machine learning work to automate the classification of leaves in studies of modern and ancient plant diversity (cf. Wilf *et al*., 2016), and could help quantify the nature and rate of leaf shape change during development (e.g. McLellan 1993) as well as investigate how leaf shape varies as a function of a tree’s aspect.

For angiosperms, “leaf size and shape are selected by climate and are strongly correlated with climatic variables” (Huff *et al*. 2003, p. 266) and a clear next step is to apply our methods to angiosperm leaves in the context of climatic and palaeoclimatic analysis (e.g. Peppe *et al*., 2011). In particular, given that our topological features measure the depth of indentations in the leaf margin, we are particularly interested to undertake quantitative analyses of angiosperm leaf margins. Such data could also feed into the Climate Leaf Analysis Multivariate Program (CLAMP) (Wolfe 1990; Spicer 2008; Yang *et al*. 2011), which uses discrete categories to describe aspects of leaf form, and may enhance characters relating to the leaf margin (such as the regularity and closeness of leaf teeth) and the overall shape of leaves.

The methods we have described could also be used to quantify other planar shapes produced by plants such as the sepals, petals, and tepals of flowers, which may enhance studies of the relationship between morphology and pollination biology (cf. Mander *et al*., 2020). As an illustration of the potential wider applicability of our methods, the long-shoot and short-shoot leaves of our modern *Ginkgo* leaves are well separated by our topological approach with minimal overlap between these two discrete classes (Fig. 5c). Given that long-shoot leaf morphology is thought to arise from the hydraulic limitation of long-shoot leaves during development (Leigh *et* al. 2011), this highlights that our methods may be usefully applied to the problem of quantifying the relationship between morphology and the underlying physiological and developmental processes that are responsible for the generation of organic form.

## Acknowledgements

We are grateful to three anonymous reviewers whose comments on a previous version of this work substantially improved the research. We are grateful to Peta Hayes for assistance with locating, accessing, and photographing fossil *Ginkgo* leaves in the collections of the Natural History Museum, London. WM acknowledges NSF grant DMS-1722995, MB was partially supported by NSF grant DMS-1953244.

## Author contributions

LM and WM designed the research, LM generated the leaf dataset, HH and MB performed experiments and analysed data with input from WM. HH, MB, WM and LM wrote the manuscript. HH and MB contributed equally.

## Supplementary Information

A dataset of modern and fossil *Ginkgo* leaf images.

## Notes

### Competing Interest Statement

The authors have declared no competing interest.

### Summary of Updates

Edits to the text following peer review.

## References

Bacon KL, Belcher CM, Haworth M, McElwain JC. 2013. Increased Atmospheric SO2 Detected from Changes in Leaf Physiognomy across the Triassic– Jurassic Boundary Interval of East Greenland. PloS ONE 8: e60614.

Barthelemy D, Caraglio Y. 2007. Plant Architecture: A Dynamic, Multilevel and Comprehensive Approach to Plant Form, Structure and Ontogeny. Annals of Botany 99: 375–407.

Bauer M, Bruveris M, Harms P, Møller-Andersen J. 2017. A numerical framework for Sobolev metrics on the space of curves. SIAM Journal on Imaging Sciences 10: 47–73.

Bauer M, Bruveris M, Charon, Møller-Andersen J. 2019. A relaxed approach for curve matching with elastic metrics. ESAIM: Control, Optimisation and Calculus of Variations 25: 72.

Bertalmio M, Sapiro G, Caselles V, Ballester C. 2000. Image inpainting. SIGGRAPH ‘00: Proceedings of the 27th Annual Conference on Computer Graphics and Interactive Techniques 417–424.

Bookstein FL. 1996. Biometrics, Biomathematics and the morphometric synthesis. Bulletin of Mathematical Biology 58: 313–365.

Bucksch A, Atta-Boateng A, Azihou AF, Battogtokh D, Baumgartner A, Binder BM, Braybrook SA, Chang C, Coneva V et al. 2017. Morphological Plant Modeling: Unleashing Geometric and Topological Potential within the Plant Sciences. Frontiers in Plant Science 8: 900.

Carvalho MA, Turgeon R, Owens T, Niklas KJ. 2017. The hydraulic architecture of Ginkgo leaves. American Journal of Botany 104: 1285–2017.

Chitwood DH, Klein LL, O’Hanlon R, Chacko S, Greg M, Kitchen C, Miller AJ, Londo JP. 2016. Latent developmental and evolutionary shapes embedded within the grapevine leaf. New Phytologist 210: 343–355.

Chitwood DH, Otoni WC. 2017. Morphometric analysis of Passiflora leaves: the relationship between landmarks of the vasculature and elliptical Fourier descriptors of the blade. Gigascience 6: giw008.

Christianson ML, Niklas KJ. 2011. Patterns of diversity in leaves from canopies of Ginkgo biloba are revealed using specific leaf area as a morphological character. American Journal of Botany 98: 1068–1076.

Critchfield WB. 1970. Shoot growth and heterophylly in Ginkgo biloba. Botanical Gazette 131: 150–162.

Dörken VM. 2013. Morphology, anatomy and vasculature in leaves of Ginkgo biloba L. (Ginkgoaceae, Ginkgoales) under functional and evolutionary aspects. Feddes Repertorium 124: 80–97.

Johnson KR. 1992. Leaf-fossil evidence for extensive floral extinction at the Cretaceous-Tertiary boundary, North Dakota, USA. Cretaceous Research 13: 91–117.

Klassen E, Srivastava A, Mio M, Joshi SH. 2004. Analysis of planar shapes using geodesic paths on shape spaces. IEEE transactions on pattern analysis and machine intelligence 26: 372–383.

Laga H, Kurtek S, Srivastava A, Miklavcic SJ. 2014. Landmark-free statistical analysis of the shape of plant leaves. Journal of theoretical biology 363: 41– 52.

Laga H, Kurtek S, Srivastava A, Golzarian M, Miklavcic SJ. 2012. A Riemannian elastic metric for shape-based plant leaf classification. 2012 International Conference on Digital Image Computing Techniques and Applications (DICTA), IEEE, 1–7.

Leaf Architecture Working Group. 1999. Manual of Leaf Architecture - morphological description and categorization of dicotyledonous and net-veined monocotyledonous angiosperms. 65p.

Leigh A, Zwieniecki MA, Rockwell FE, Boyce CK, Nicotra AB, Holbrook NM. 2011. Structural and physiological correlates of heterophylly in Ginkgo biloba L. New Phytologist 189: 459–470.

Leigh EG. Jr. 1999. Tropical Forest Ecology. Oxford University Press, New York.

Li M, Duncan K, Topp CN, Chitwood DH. 2017. Persistent homology and the branching topologies of plants. American Journal of Botany 104: 349–353.

Li M, Frank MH, Coneva V, Mio W, Chitwood DH, Topp CN. 2018a. The persistent homology mathematical framework provides enhanced genotype-to-phenotype associations for plant morphology. Plant Physiology 177: 1382–1395.

Li M, An H, Angelovici R, Bagaza C, Batushansky A, Clark L, Coneva V, Donoghue M, Edwards E et al. 2018b. Topological data analysis as a morphometric method: using persistent homology to demarcate a leaf morphospace. Frontiers in Plant Science 9: 553.

Lin S, Niklas KJ, Holscher D, Hui C, Ding Y, Shi P. 2020. Leaf shape influences the scaling of leaf dry mass vs. area: a test case using bamboos. Annals of Forest Science 77: 11.

McLellan, T. 1993. The roles of heterochrony and heteroblasty in the diversification of leaf shapes in Begonia dreigei (Begoniaceae). American Journal of Botany 80: 796–804.

Mander L, Parins-Fukuchi C, Dick CW, Punyasena SW, Jaramillo C. 2020. Phylogenetic and ecological correlates of pollen morphological diversity in a Neotropical rainforest. Biotropica 53: 74–85.

McElwain JC, Steinthorsdottir M. 2017. Paleoecology, ploidy, paleoatmospheric composition, and developmental biology: a review of the multiple uses of fossil stomata. Plant Physiology 174: 650–664.

McLellan T. 1993. The Roles of Heterochrony and Heteroblasty in the Diversification of Leaf Shapes in Begonia dregei (Begoniaceae). American Journal of Botany, 80: 796–804.

Nicotra AB, Leigh A, Boyce CK, Jones CS, Niklas KJ, Royer DL, Tsukaya H. 2011. The evolution and functional significance of leaf shape in the angiosperms. Functional Plant Biology 38: 535–552.

Niklas KJ, Christianson ML. 2011. Differences in the scaling of area and mass of Ginkgo biloba (Ginkgoaceae) leaves and their relevance to the study of specific leaf area. American Journal of Botany 98: 1381–1386.

Otter N, Porter MA, Tillmann U, Grindrod P, Harington HA. 2017. A roadmap for the computation of persistent homology. EPJ Data Science 6: 17.

Peppe DJ, Royer DL, Cariglino B, Oliver SY, Newman S, Leight E, Enikolopov G, Fernandez–Burgos M, Herrera F et al. 2011. Sensitivity of leaf size and shape to climate: global patterns and paleoclimatic applications. New Phytologist 190: 724–739.

Royer DL, Wilf P, Janesko DA, Kowalski EA, Dilcher DL. 2005. Correlations of climate and plant ecology to leaf size and shape: Potential proxies for the fossil record. American Journal of Botany 92: 1141–1152.

Royer D, McElwain J, Adams J. 2008. Sensitivity of leaf size and shape to climate within Acer rubrum and Quercus kelloggii. New Phytologist 179: 808–817.

Royer DL, Meyerson LA, Robertson KM, Adams JM. 2009. Phenotypic plasticity of leaf shape along a temperature gradient in Acer rubrum. PloS ONE 4: e7653.

Shi P, Li YR, Hui C, Ratkowsky DA, Yu XJ, Niinemets Ü. 2020. Does the law of diminishing returns in leaf scaling apply to vines? – Evidence from 12 species of climbing plants. Global Ecology and Conservation 21: e00830.

Spicer RA. 2008. CLAMP. In:V. Gornitz (Ed.), Encyclopedia of Paleoclimatology and Ancient Environments. Springer, Dordrecht, pp. 156–158.

Sun B, Dilcher DL, Beerling DJ, Zhang C, Yan D, Kowalski E. 2003. Variation in Ginkgo biloba L. leaf characters across a climatic gradient in China. Proceeding of the national Academy of Sciences, USA 100: 7141–7146.

Thompson, D’AW. 1942. On Growth and Form. Second Edition, Cambridge University Press, Cambridge.

Webster M, Sheets HD. 2010. A practical introduction to landmark-based morphometrics. Paleontological Society Special Papers 16: 163–188.

Weight C, Parnham D, Waites R. 2008. LeafAnalyser: a computational method for rapid and large-scale analyses of leaf shape variation. Plant Journal 53: 578–586.

Wilf P, Zhang S, Chikkerur S, Little SA, Wing SL, Serre T. 2016. Computer vision cracks the leaf code. Proceeding of the national Academy of Sciences, USA 113: 3305–3310.

Wolfe JA. 1990. Palaeobotanical evidence for a marked temperature increase following the Cretaceous/Tertiary boundary. Nature 343: 153–156

Xie S, Sun B, Yan D, Du B. 2009. Altitudinal variation in Ginkgo leaf characters: Clues to paleoelevation reconstruction. Science in China Series D: Earth Sciences 52: 2040–2046.

Yang J, Spicer RA, Spicer TEV, Li C-S. 2011. ‘CLAMP’ Online: a new web-based palaeoclimate tool and its application to the terrestrial Paleogene and Neogene of North America. Palaeobiodiversity and Palaeoenvironments 91: 163–183.

